# Virus-host protein co-expression networks reveal temporal organization and strategies of viral infection

**DOI:** 10.1101/2023.10.10.561729

**Authors:** Jacobo Aguirre, Raúl Guantes

**Affiliations:** Centro de Astrobiología (CAB), CSIC-INTA, ctra. de Ajalvir km 4, 28850 Torrejón de Ardoz, Madrid, Spain; Grupo Interdisciplinar de Sistemas Complejos (GISC), Madrid, Spain; Department of Condensed Matter Physics and Material Science Institute ‘Nicolás Cabrera’, Science Faculty, Universidad Autónoma de Madrid, Madrid, Spain; Condensed Matter Physics Center (IFIMAC), Science Faculty, Universidad Autónoma de Madrid, Madrid, Spain

**Author notes:** Corresponding authors: Raúl Guantes, Jacobo Aguirre.

## Abstract

Viral replication is a complex dynamical process involving the global remodelling of the host cellular machinery across several stages. In this study, we provide a unified view of the virus-host interaction at the proteome level reconstructing protein co-expression networks from quantitative temporal data of four large DNA viruses. We take advantage of a formal framework, the theory of interacting networks, to describe the viral infection as a dynamical system taking place on a network of networks where perturbations induced by viral proteins spread to hijack the host proteome for the virus benefit. Our methodology demonstrates how the viral replication cycle can be effectively examined as a complex interaction between protein networks, providing useful insights into the viral and host’s temporal organization and strategies, key protein nodes targeted by the virus and dynamical bottlenecks during the course of the infection.

## INTRODUCTION

Throughout an infection, viruses hijack different cellular processes of the host cell, dynamically remodelling the abundance, modifications and interactions of many proteins. Recent improvements in proteomic techniques have enabled precise quantification of thousands of host and viral proteins at different times post-infection^1^. By simultaneously measuring the abundance of host and viral proteins in a reference (non-infected) situation, these experiments open the possibility to examine which proteins are significantly altered during the infection and when, thus highlighting biologically relevant pathways and potential therapeutic targets^1–3^.

Currently, most analyses of time-resolved proteome abundances have focused on clustering proteins by similarity in expression profiles, followed by functional analysis such as pathway or gene set enrichment analysis (GSEA)^1,4^. These studies have revealed host signalling and metabolic pathways affected by the viral infection^2,5,6^ and unveiled viral strategies to evade the immune response of the host cell^3,7–9^. On the other hand, many works have addressed the characterization of virus-host molecular interactions from a network perspective, reconstructing the protein interaction networks (PINs) for different pathogens^10–18^. This work has uncovered common structural trends in virus-host PINs^15,18^, such as that viral proteins tend to interact with host protein hubs^12,13^. Despite the usefulness of these studies to comparatively establish principles of viral infection mechanisms^10,16^, the nodes of these networks include only proteins with direct physical interactions, thus excluding all indirect regulatory interactions such as transcriptional regulation. Furthermore, and especially for large DNA viruses, the progression of the viral infection depends on the establishment of temporally tuned virus-host and virus-virus protein interactions^19^. With few exceptions^20–22^, most virus-host PINs provide only a static picture of the molecular interactome. In summary, these approaches do not take full advantage of the powerful tools recently developed in the framework of the analysis of dynamical processes on networks in interaction^23,24^.

In this work, we aim to bridge the gap between the systems level perspective of the viral infection process and its dynamics, by reconstructing and investigating topological and dynamical features of virus-host protein co-expression networks. We make use of the theory of interacting networks^25^, a mathematical framework designed to analyse the dynamical processes that take place on networks of networks^24^ that is, complex networks connected through a limited number of interlinks, both in equilibrium^26^ and out-of-equilibrium^27^ environments. In this methodology, network spectral properties (i.e. the eigenvalues and eigenvectors of the adjacency matrices associated with the networks under study^28^) become critical tools to describe the systems’ complex dynamics, and will help us to perform a thorough comparative analysis of the co-expression networks of four large DNA viruses, with different replication cycle durations and mechanisms. We will focus on (i) how the global network structure reflects the division between virus and host temporal organization, revealing a modular rearrangement of the host proteome during the course of the infection, (ii) how dynamics is naturally included in the network structure through the response time of each node, linking topological features to the temporal program triggered by the viral infection to remodel the host cell machinery, and (iii) how the analysis of node importance provides valuable information on key host proteins and biological processes targeted by the virus. Overall, the multidisciplinary approach followed in this work offers a novel perspective on the global temporal organization of a viral replication cycle, and can provide useful biological insight on relevant players and mechanisms of viral infection up to now overlooked in more standard analytical approaches.

## 2. RESULTS

### 2.1 Virus-host protein dynamics described as a network of networks

To provide a global picture of how viral infection impacts the host proteome, we construct co-expression networks from quantitative temporal proteomics data of four large DNA viruses during their replication cycle inside a host: HSV-1, EBV, HCMV and VACV (Fig. 1, see *Methods* for full details on their construction, and Table S1 for information on the viruses and data sources analysed in this work). In these networks, each node represents a protein that has been differentially expressed with respect to the non-infected case at least at one time, and two nodes are connected through a link if their expression patterns change proportionally (or reciprocally) across time (*Methods*). One interesting property of these networks is their modularity, a measure of the strength of division of a network into communities. All virus-host co-expression networks show a modular topology (modularity coefficient > 0.6, Table S2), that makes them behave as networks of networks^23,24^. In Fig. 1 (right panels) we plot the community network for each virus-host system, that is, a simplified representation of the co-expression networks where each node corresponds to a full community and links connect communities that are interconnected in the whole networks (see Table S3 for the topological description of the different communities).

**Figure 1.**
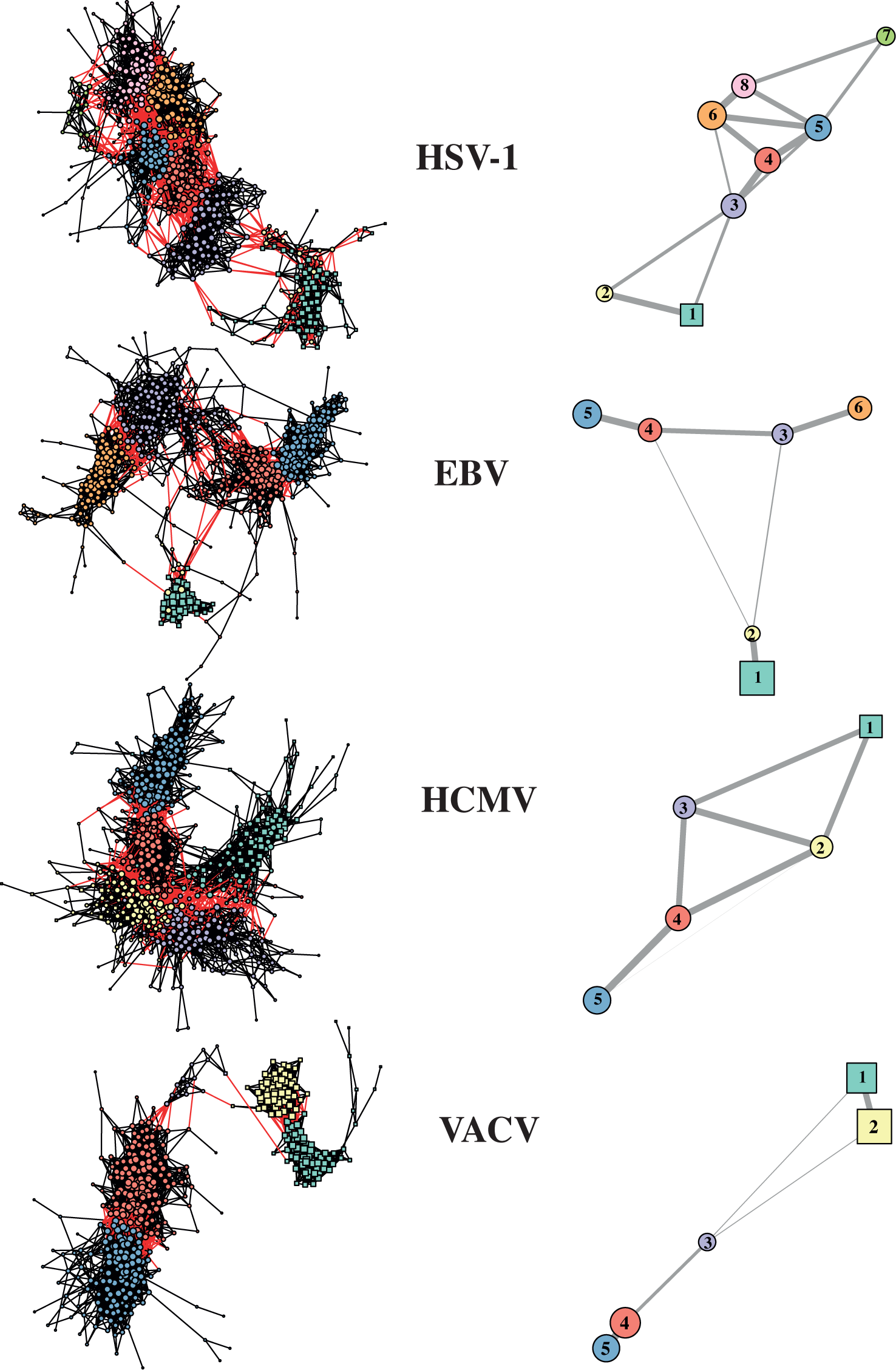
Virus-host protein co-expression networks are modular. *Left panels*: Virus-host co-expression networks of four different viruses (Tables S1 and S2). Circles: host proteins. Squares: Viral proteins. Node size is proportional to node degree, node colour denotes community membership and red links are inter-community connections. *Right panels:* Network of communities for each virus, highlighting the modular structure of the co-expression network. Colour codes as in left panels. Viral nodes form separate communities (squares) connected to the host network (circles) by one or two small communities, except for the HCMV. Node size is proportional to average internal degree of each community, and edges are weighted with the logarithm of the number of inter-community links.

Importantly, it is evident from Fig. 1 that in all co-expression networks virus and host proteins form clearly differentiated subnetworks. In three of them, the activities of viral proteins are so synchronized that they form a single community (two in VACV) of large average degree, i.e., high internal connectivity. In these systems, the virus-host connection takes place through one or two host communities of low size and average degree (Fig. 1 and Fig. S3), acting as a weak, but fundamental bridge with the rest of the host network. These *connector communities* are number 2 in HSV-1 and EBV and 3 in VACV (Fig. 1 right panels), and they are formed by around 10-20 peripheral host nodes weakly connected among them. Finally, communities are interconnected through *connector nodes* (i.e., those connected to other communities) of a wide range of degree values, but with average degree significantly larger than the *internal nodes* (i.e., nodes that only show connections to their own community), as shown in Fig. S3. In summary, the topology of the co-expression networks obtained for the infection dynamics of the four viruses under study can be described as a modular structure where virus proteins are clustered in strongly internally-connected communities. In three of the four viruses, these viral communities are clearly differentiated from the host proteins via small host connector communities. Moreover, the different host communities are interconnected through proteins of relatively large degree, that is, through densely connected nodes.

### 2.2 Network topology reflects response dynamics

The co-expression networks associated with the virus-host interaction described here are static, but reflect reliable information about the dynamical process associated with each viral replication cycle. To see this, we assigned to every protein a single response time as the first time to exceed the cut-off in fold-change abundance (*Methods* and Table S2). Figure 2 (left panels) shows the correspondence between the response time of every protein and its network distance (length of shortest path) to the viral immediate early genes (IEGs, Table S1). In HSV-1, EBV and VACV, the viral proteins are the ones to be firstly activated, and a “response wave” spreads over the network, starting on the viral communities and advancing toward the furthest host proteins when time increases. Note that the dispersion in the EBV has reached most of the host at 24h, and therefore the correlation between response time and the distance to the firstly activated proteins is still detectable but less steep (a finer temporal resolution for EBV would be needed to improve these results). In contrast to the other viruses, the HCMV viral proteins (pale blue squares in Fig. 2) show a slow response pattern (≥ 48h for most of them). This behaviour results in a two-phase complex pattern of dispersion over the network, one starting in the first steps of the infection, and the second one beginning around the outburst at 48h. This pattern reflects the actual life cycle of the HCMV, which is long and dynamically very complex, involving both nuclear and cytoplasmic phases^29,30^.

**Figure 2.**
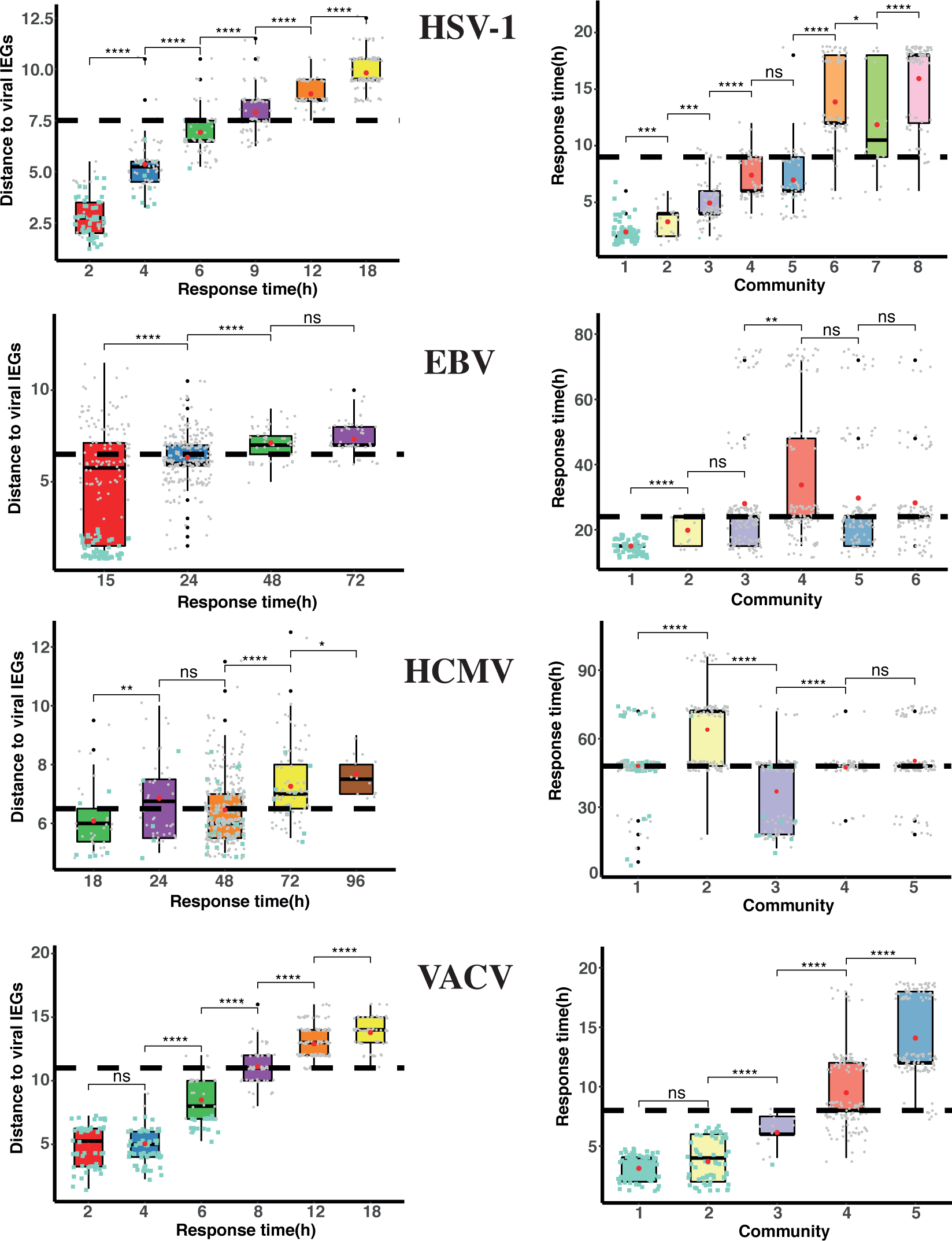
Protein response dynamics at the protein co-expression networks. *Left panels*: Distributions (shown as boxplots, red dots are distribution means) of average network distances to viral immediate early genes (IEGs, Table S1). Response times are defined as the first experimental time point in which protein abundance exceeds the cut-off in fold-change (*Methods* and Table S2). *Right panels:* Distributions (boxplots) of protein response times for the different communities in which each co-expression network is divided.

The detection of response waves over the networks enables us to analyse the virus-host interaction as a dynamical system taking place on the co-expression network, where relevant perturbations created by the viral machinery spread over the host cells in an organized manner. The right panels in Fig. 2 show the relationship between response times and the different communities in which each network is divided. In HSV-1 and VACV the correspondence is clear, implying that the perturbation caused by the infection in the host proteome spreads sequentially over the different communities. Furthermore, note that in these two cases the spreading follows strictly the connection patterns among communities (Figs. 1 and 2, right panels). Regarding EBV, and in agreement with the response time-distance correlation studied above, the perturbation has reached most communities in the first stages of the infection, and only a slightly later response of community 4 is detected. Finally, the HCMV network is more difficult to analyse due to the extensive cross-talk between viral and host proteins^31–33^, resulting in strong connectivity between communities (Fig. 1 right panel) and slow response dynamics of viral and host nodes. Community 3 responds sooner on average, likely because it contains several early rising viral proteins driving faster changes in the host, which spread later to communities 4 and 5. Community 1, on the other hand, contains many late-rising viral proteins activating the host community 2 in a second stage of the infection process (Fig. 2).

### 2.3 The theory of interacting networks unveils infection complex dynamics

The co-expression networks that we are studying here can be considered as networks of networks, where the virus communities interact with very modular co-expression host networks through a relatively small number of interlinks. While the links in each of these networks represent a complex mixture of biochemical and gene regulatory interactions, many of them unknown, and not a clearly defined process spreading on a physical structure (as it could happen in the dispersion of an illness on two populations that get in contact, for example), the systems can be described as response waves that start in the viral communities and sequentially spread over the whole host networks.

Extensive work has been developed on the last two decades to describe the nature of complex interactions in networks of networks and the processes that take place on them, and today it is clear that some of their properties drastically differ from those of dynamical systems spreading on isolated networks, as it arose studying network robustness^34^ or synchronization^35,36^, to cite just a few examples. The theory of interacting networks^25–27^ (TIN from now on) approaches the analysis of dynamical processes evolving on networks of networks as a competition for a topological measure called eigenvector centrality, and yields that the network spectral properties of the networks under study determine the systems’ complex dynamics (see *Methods*). The maximum eigenvalue of the adjacency matrix λ_1_ is a proxy for the strength of a network, and its associated eigenvector has been used in many different contexts with the name of eigenvector centrality to measure the dynamical importance of nodes in a network (or a whole network in a network of networks if we add up the centrality of each network’s nodes)^28^. From this perspective, dynamical processes on interconnected networks are analysed in TIN as a competition between networks for eigenvector centrality representing importance (or influence, resources, wealth, or whatever the process is describing). The main conclusions of this theory for out-of-equilibrium systems are (i) the system’s dynamics depends (almost) exclusively on the maximum eigenvalues of the different interconnected networks and the centrality of the connector nodes, and (ii) the dynamics is fast when networks are strongly interconnected, especially if this is through connector nodes of large internal eigenvector centrality, while few and peripheral inter-network links act as bottlenecks and drastically slow down the dynamics taking place on the system.

Let us analyse how the theory of interacting networks can be used to describe the dynamics of the virus-host interaction. Table S3 presents the spectral properties of the adjacency matrix associated with the virus network, the host network, and each of the communities in which the co-expression networks have been divided. The distribution of eigenvector centrality of internal nodes (nodes connected only to other members of the same community) and connector nodes (nodes connecting to other communities) of each community in Fig. 3 shows that host communities are interconnected through nodes that, on average, have larger centrality than their internal nodes. From the perspective of TIN, the connection patterns of HSV1, EBV and VACV suggest a dynamic view in which the perturbation originated by the virus first affects a small number of proteins (the connector communities), acting as dynamic bottlenecks between the virus and the rest of the host. In a second stage, and due to the interconnection between communities through important nodes, the perturbation spreads easily throughout the rest of the host. Furthermore, the drastic imbalance between the centrality accumulated in the virus and the host network (Table S3) is a proof of the isolation of the virus network with respect to the host, while the much more balanced centrality spreading among the different communities of the host networks shows that, while they are clearly topologically differentiated, are nonetheless efficiently interconnected. Note, finally, that in the case of EBV, the viral proteins are so strongly internally connected that its spectral gap (the difference between the two largest eigenvalues, *Λ*_1_–*Λ*_2_) is much larger than the rest of viral networks (a typical property of cliques, or networks where every two nodes are linked). This is due to the fact that the EBV viral proteins behave as an early activated, influential, and highly synchronized cluster.

**Figure 3.**
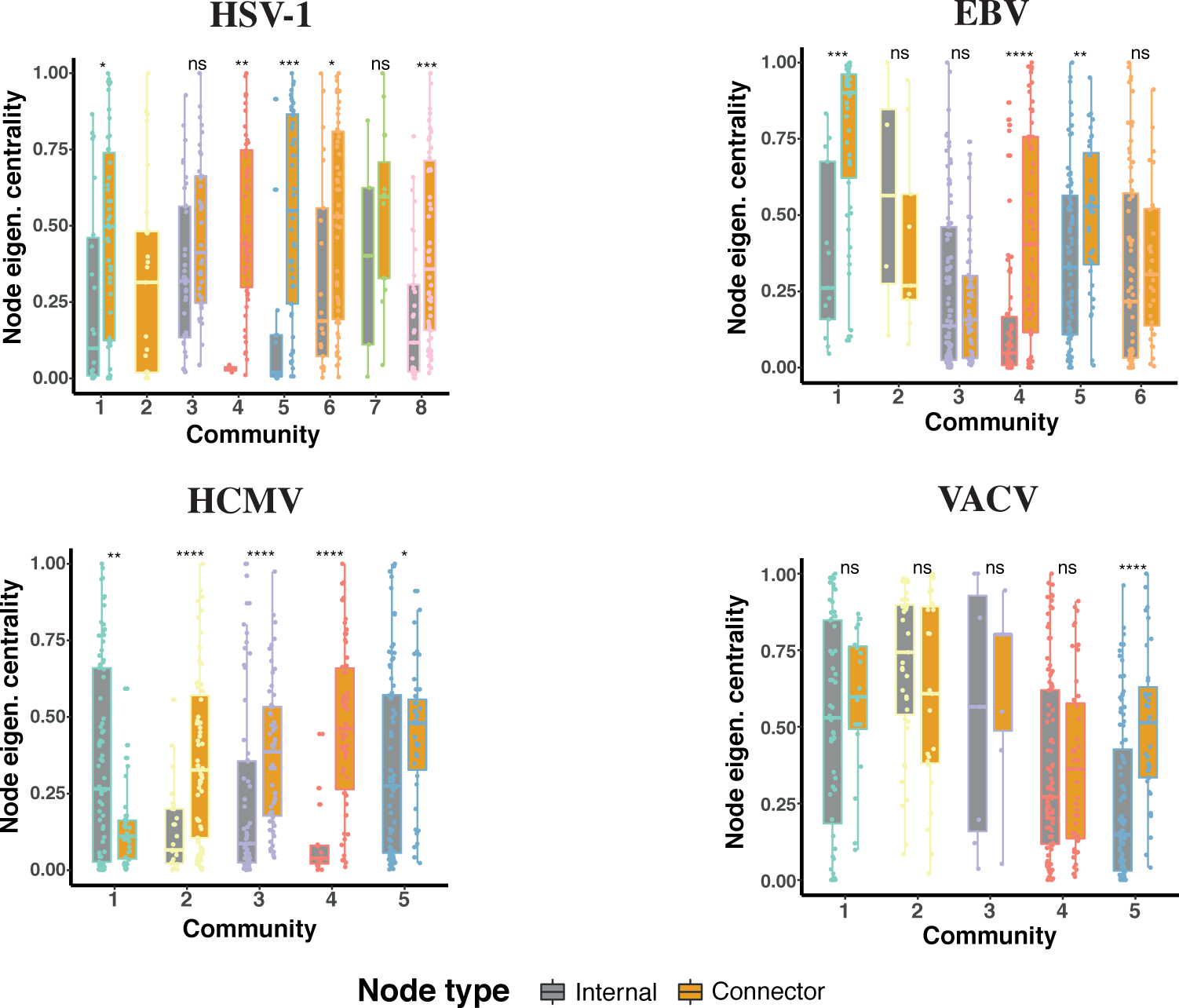
Eigenvector centrality of internal nodes and connector nodes. Distributions of node eigenvector centralities for each virus type and community, separating internal nodes (nodes connected only to other members of the same community, grey boxes) from connector nodes (nodes connecting to other communities, orange boxes). Significance p-values for difference of the mean between internal and connector nodes are calculated with Wilcoxon test (ns: p>0.05; *: p ≤0.05, **: p ≤0.01; *** : p ≤0.001; **** : p ≤0.0001).

### 2.4 Node centrality reveals functional structures

Degree centrality measures the importance of a node in a network based on the number of its neighbours, and therefore represents strictly local influence. Eigenvector centrality, on the contrary, measures the importance of a node based on the relevance of such neighbours, and in practice describes the power of a node to affect other nodes in the whole network. Usual measures of modularity that only take into account the topological structure of networks (based on node degree), if applied to networks where dynamics take place, could yield an incomplete description of the system. On the other hand, using eigenvector centrality to describe the structure of the network can provide a community partition that reveals important dynamical features^37^.

Following this idea, we show in Fig. 4 the node degree versus node eigenvector centrality for each virus, with colours representing the community membership for each protein. This combination of the local and global influence of each protein/node allows us to check, regarding the node clusters present in the plot, whether the communities associated with the system and plotted in Fig. 1 show some hidden dynamical structures, and detect the most relevant nodes in the infection process. Figure 4 shows a hierarchical structure of “tongues” for the virus-host networks, in which the viral community is neatly separated from the rest, and connected by a small set of nodes (with the exception of HCMV) to the larger host cluster. The host communities shown in Fig. 1 appear as successive layers in the degree/eigencentrality plots. In the case of EBV, for instance, two subclusters are clearly distinguished, corresponding to communities 3/6 and 4/5 (see also Fig. 1 left).

**Figure 4.**
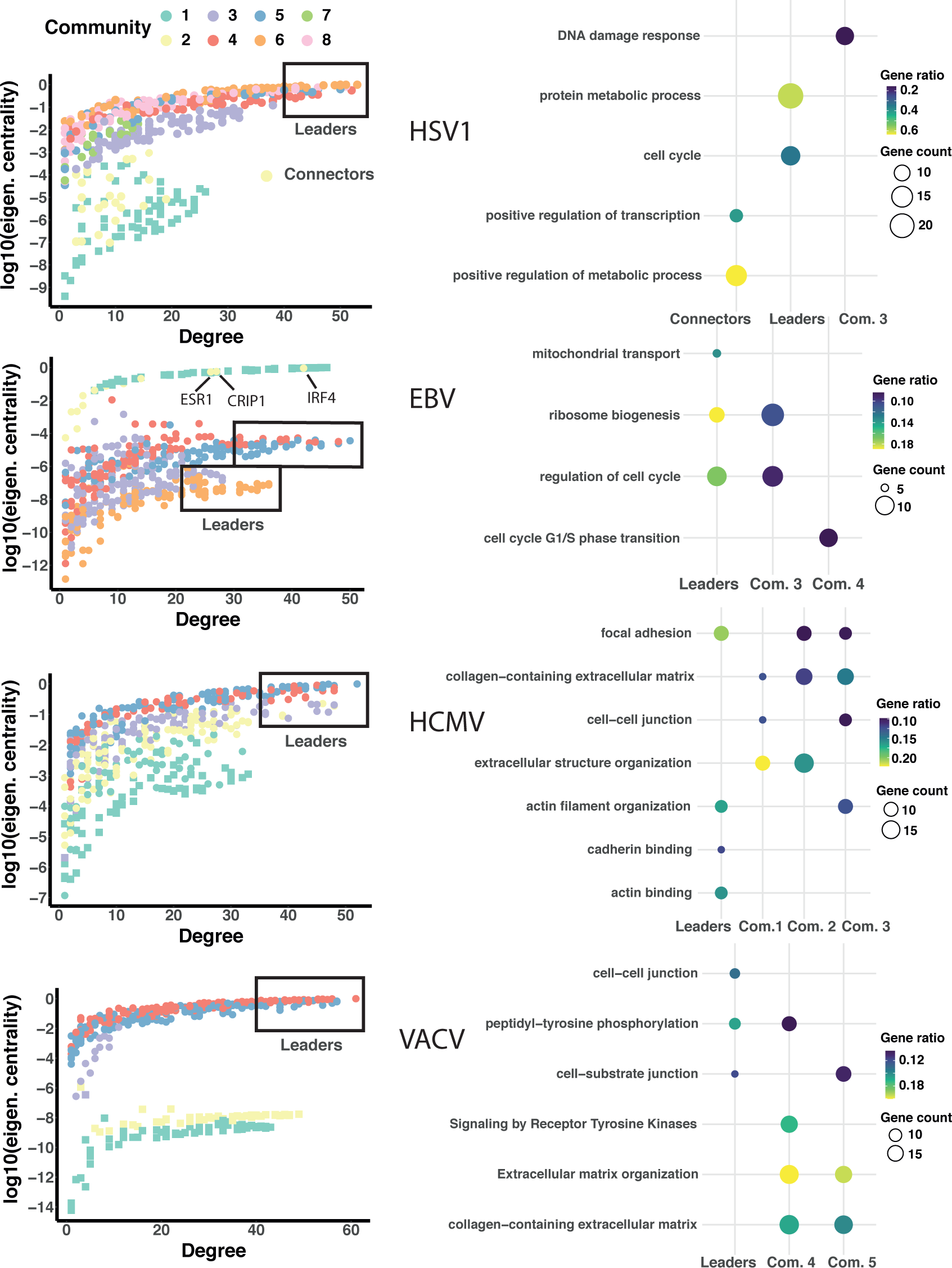
Node centrality and functionality in community structure. *Left panels:* Degree versus eigenvector centrality of network nodes. Colours correspond to communities as in Figs. 1-3. Square symbols are viral proteins, circles are host proteins. *Right panels*: Functional analysis of leader nodes, virus-host connector nodes and communities highlighting the most significant biological processes hijacked by the different viruses (see *Methods*).

Next, we explore whether the community and dynamical partitions in Fig. 4 reveal preferential functional roles within the host proteome, performing over-representation analysis (ORA) of biological functions (*Methods*). We focus on communities (especially virus-host connector communities) and on *leader proteins*: those that combine large degree (hubs) and large eigenvector centrality (central nodes), enclosed within black boxes in Fig. 4. In the following we discuss the main findings in each of the virus-host systems studied (Fig. 4 right) and relate them to the known biology of the particular viral infection process.

#### HSV1

One salient topological feature of the HSV1 virus-host network is the presence of a sizeable community of connector nodes between the viral and host subnetworks (Fig. 4 left). Functional analysis shows that ∼65% of the proteins in this community (all downregulated) are related to positive regulation of cellular metabolism (Fig. 4 right), and many of them are transcription factors involved in the regulation of metabolic processes and protein degradation (Table S4). One key mechanism used by HSV1 to escape immunity and promote productive infection is to hijack the ubiquitin-proteasome system^38,39^, the main pathway for degradation and functional modification of proteins, orchestrated by the immediate early viral gene *ICP0*^40^. Notably, the connector community contains the proteins most sharply downregulated after viral replication induction^41^ (Fig. S4A, yellow time series). Several of these connector proteins, involved in the ubiquitin-proteasome pathway, are known to be targeted for degradation by *ICP0*^40^ (Table S4).

Leader proteins (those with high degree and eigenvector centrality, Fig. 4 left) play significant roles in the regulation of the cell cycle and in protein degradation (Fig. 4 right and Table S4). *ICP0* is known to dysregulate cell cycle, inducing arrest to enhance viral replication^42^. The alteration of the ubiquitin-proteasome system is also significantly detected in some communities, especially in community 3 and the smaller community 7 (Table S4). Community 3 is also rich in proteins of the DNA damage response pathway (Fig. 4 right and Table S4). Again, this pathway is inhibited upon HSV1 infection to favour virus survival, mainly in an *ICP0* dependent manner^39,40^.

#### EBV

The EBV co-expression network has a small set of virus-host connector nodes (all assigned to community 2, Fig. 1). Several upregulated proteins of this community are associated with the complement system^5^, whose alteration may be connected to the tight association between EBV infection and multiple sclerosis^43,44^. The most salient topological feature of the connector community, however, is the presence of three host proteins densely connected to the viral subnetwork (Fig. 4 left): Interferon Regulatory Factor 4 (IRF4), which is strongly upregulated, CRIP1 and ESR1. IRF4 has been shown to facilitate EBV lytic reactivation^45^ and is implicated in the differentiation of B cells toward plasma cells associated with many human lymphoid malignancies^46^. CRIP1, on the other hand, is strongly downregulated and has been identified as a biomarker of EBV-associated nasopharyngeal carcinoma^47^. The estrogen receptor-1 protein ESR1 (upregulated), is known to induce the expression of the immediate early viral gene *BZLF1* and has been also associated to poor prognosis of nasopharyngeal carcinoma^48^.

A second distinctive topological feature of the EBV network is the existence of two clusters of communities (communities 4/5 and 3/6, Fig. 1) both connected to the viral subnetwork through the connector community 2. This structure is apparent in the degree-eigencentrality plots, Fig. 4 left, as two ‘tongues’ containing the most connected and central nodes of both clusters. The leader proteins of communities 3/6 are enriched in proteins involved in ribosome biogenesis, all downregulated. The translational machinery of the host cell is modified by different viruses to selectively favour viral replication, and EBV is known to interfere with ribosomal host proteins to activate specific viral genes^49^. Leaders of communities 4/5 contain especially proteins involved in cell cycle regulation. Like other herpes viruses, EBV induces cell cycle arrest at G1/S-phase to promote viral replication^50^. Signatures of this viral reprogramming are significant and noticeably in communities 3 and 4, which connect the two large clusters (Fig. 4 right and Table S4).

#### HCMV

The life cycle of HCMV is dynamically very complex, with different groups of viral proteins temporally controlled to be expressed at specific phases of replication^31,51^. Time dependent transcriptome and proteome analyses have revealed up to seven different temporal classes^33,52^, with some viral genes expressed as late as 3-4 days after the onset of productive infection. This complexity is apparent in the virus-host co-expression network, since viral proteins do not form a clearly independent community from the host as in the other viruses (Fig. 4 and Table S3), highlighting the intricate modulation of the host cell proteome during the course of HCMV replication^29^.

The most prominent biological feature throughout the network is a large and significant hallmark of altered host genes involved in focal adhesion as well as in the reorganization of the extracellular matrix and the cytoskeleton. Leader proteins are all downregulated, and mostly involved in focal adhesion and actin filament organization (mainly through cadherin and acting binding, Fig. 4 right). Notably, one of the most connected proteins is adducin 3 (ADD3), involved in the assembly of the actin filament network at sites of cell-cell contact. This gene is a marker of biliary atresia^53^, a pathology also associated to HCMV infection^54^. Functions related to adhesion and extracellular matrix organization are also over-represented in the rest of the communities, consistent with a global downregulation of these processes in HCMV infected cells^55^. The signatures observed here are also in agreement with a general reprogramming of mesenchymal-to-epithelial (MET) and epithelial-to-mesenchymal (EMT) transition pathways recently uncovered in HCMV infection^56^. Dysregulation of these pathways, that are associated to cancer progression and prognosis^57^, is temporally organized across different communities (Table S4).

#### VACV

Poxviruses are characterized by conducting the whole infectious cycle within the cytoplasm of the host cell, without direct involvement of the nucleus. As an archetype of this viral family, vaccinia encodes virtually all the proteins necessary for replication in the cytoplasm, which takes place in ‘replication factories’ that compartmentalize and help to protect the viral genome from immune detection. The low interconnectedness observed between viral and host communities for VACV likely reflects this autonomous replication mechanism. The distinctive feature of the VACV co-expression network is the existence of two separated viral communities (communities 1 and 2, Fig. 1), corresponding to two well differentiated time profiles (Fig. S4D, green and yellow time series). Community 1, which includes the viral nodes most sharply activated upon infection, contains the proteins initially involved in the core replication machinery^58^ (Table S4). On the other hand, the nodes belonging to the viral community 2 activate in a more graded manner and at later times. This community includes the proteins involved in different viral phases, from immature virion assembly to envelope formation^59^, Table S4. The viral subnetwork partition thus reflects the dynamic organization of the VACV life cycle.

There are around 50 leader proteins, all downregulated, belonging to communities 4 and 5, Fig. 4 left. These include many plasma membrane proteins, which are globally modulated during VACV infection to evade the immune system and alter the cell host motility^60^. Of note, many of these are proteins involved in the Ephrin signalling pathway^61^ (Table S4) which is used by different viruses for viral entry^62^. Ephrins are known to modulate cell adhesion and migration properties^61,63^. The strong downregulation of Ephrin receptors observed within the leader community, which could facilitate cell-cell repulsion and increase cell migration^63^, suggests the mechanism employed by VACV to induce cell motility and enhance the spread of infection^64^. In agreement with this, we observed a large and highly significant functional role of both receptor tyrosine-kinase mediated signalling (involved in cell adhesion and migration) and extracellular matrix re-organization in the host communities (Fig. 4 right).

In summary, the combined topological and functional analyses of our reconstructed co-expression networks highlight the key mechanisms by which different viruses replicate and modulate the host proteome to evade immunity and enhance the spreading of infection. Communities and highly central (leader) proteins preferentially reflect the alteration of a particular pathway and the organization of its temporal response.

## 3. DISCUSSION

Taking advantage of a complex network perspective, we presented here a thorough systems level analysis of the temporal program of virus and host proteome reorganization after productive viral infection, using quantitative time course data of four large DNA viruses with complex replication cycles. Despite the specific characteristics of the different viruses, the application of the theory of interacting networks (TIN) shows how the infection process can be seen as the interaction between two co-expression networks (the virus and host proteomes) in which the perturbation induced by the virus propagates in a temporally coordinated manner through the host network. Upon lytic activation, HSV1, EBV and VACV replicate in a relatively autonomous and synchronized manner independent of the dynamics of the host response. This is especially clear in the EBV co-expression network, and consistent with the fact that this virus encodes its own machinery to initiate DNA amplification and proceed to late gene expression^65^. Our reconstructed networks for these viruses show that viral proteins are connected to a small set of host proteins of low average degree. This is in contrast to the observed topology in virus-host protein interaction networks, in which viral proteins preferentially target host proteins that are densely connected (hubs)^12,13,15^. Studies in virus-host PINs suggest that host proteins targeted by viruses are evolutionarily conserved^12,66^ and can control a wide range of functions. In our case, the weak connections between viral and host subnetworks are the hallmark of a dynamical bottleneck between the onset of viral replication and the propagation of the perturbation to the host proteome. Host connector nodes may functionally represent proteins targeted by viral early genes to establish a host environment favourable for replication, as is the case in HSV1^40^ or promoting lytic activation, as in EBV infected B cells^45^.

Based on TIN’s predictions, the modularity and topological structure of the host network (where connector nodes between communities have a larger eigenvector centrality on average) suggest a second dynamical phase in which the perturbation produced in a few host proteins propagates more efficiently between communities. Functional analyses of different communities show, in fact, that they do not act as independent functional modules but may be part of the same host pathways remodelled through different temporal stages. For instance, two or more interconnected communities are involved in the dysregulation of cell adhesion and mobility properties of the infected cells both in VACV and HCMV. Additional substructure within communities may be revealed by a joint analysis of degree/eigenvector centrality, highlighting groups of important nodes (leader proteins) with relevant functional roles. Since these are proteins that co-vary in time in a concerted way with many other proteins, they can be naturally seen as nodes synchronizing the activity of other nodes involved in similar functions.

The global structural and dynamical analysis of co-expression networks of different virus-host systems as performed here is also useful to pinpoint common and specific mechanisms of viral immune evasion and spreading. For instance, the suppression of the SMC5/SMC6 complex both in HSV1 and EBV infection points towards an important node of the host defence mechanism^67^. Many of the proteins targeted for downregulation by VACV involved in cell adhesion and motility were also downregulated in HCMV infection^3,60^(Table S4), revealing similar strategies of these two large viruses to spread infection despite the different life cycles and mechanisms used for replication.

The co-expression networks reconstructed here are based on a parsimonious procedure and a sensible measure of proportionality. While different methods exist to infer co-expression networks, mainly in the context of gene transcription and many experimental samples^68–70^, our procedure recovered a connected virus-host network dynamically meaningful. The correspondence between dynamics and network topology was especially clear in the two viruses with shorter life cycles and better temporal resolution, HSV1 and VACV. These results show that the methods used here, combined with the theory of interacting networks, can become useful analysis tools as quantitative temporal viromics increases its multiplexing capacity improving temporal resolution.

## 4. METHODS

### 4.1 Virus-host temporal proteomic data

We gathered experimental protein abundances from time resolved proteome experiments employing tandem mass tags (TMM) and triple-stage mass spectrometry (MS3), which considerably reduce experimental noise and allow precise quantitation of thousands of viral and host proteins^1^. In particular, we analysed protein abundance data from four large double-stranded DNA viruses. Three of them belong to the herpes family: Herpes Simplex Virus 1(HSV-1)^41^, Epstein-Barr Virus(EBV)^5^ and Human Cytomegalovirus (HCMV)^33^. The fourth virus analysed, Vaccinia virus (VACV)^3^ is an archetype of the pox viral family and genetically related to variola virus, responsible for smallpox. Although up to 19 different viruses have been studied with temporal proteomic techniques^1^, we selected those viruses with a relatively large genome and a sizable number of quantified viral proteins, and at least five experimental time points. Details on the viruses and data sets are provided in Supplemental Table S1.

Original data expressed in pseudo-counts were re-analysed for differential expression with respect to a control (non-infected or uninduced samples) using the *edgeR* pipeline^71^. All viruses analysed, with the exception of HSV-1, had at least two biological replicates. We used a paired design (mock versus infected samples at each time point) controlling for differences between biological replicates, by fitting a negative binomial generalized log-linear model to protein pseudo-counts (functions *glmQLFit* and *glmQLFTest* in *edgeR*). For samples/time points without biological replicates, we estimated the significance in differential expression using dispersions from the closest time points. We found that only ∼12-18% of all quantified host proteins were differentially expressed (FDR < 0.05) with respect to the non-infected case at least at one time point, but nearly all quantified viral proteins over-expressed with respect to the control (Table S1). We note that the experimental protein quantification protocol is the same for all viruses analysed^1^, and all time-point samples per virus are analysed in the same LC-MS/MS run, which reduces experimental variability and allows for reliable comparisons across conditions (post-infection time). Nevertheless, to take into account possible effects of composition bias between samples, we used trimmed mean of M-values (TMM) normalization^72^ of data for further analyses, as has been done in other quantitative temporal viromics studies^2^.

### 4.2 Reconstruction of virus-host protein co-expression networks

Our goal is to provide a global picture of the most important players and their dynamical relationships in the virus-host infection process. We thus seek to reconstruct a manageable virus-host protein co-expression network in which the nodes are the virus/host proteins most significantly altered during the course of infection (with respect to the non-infected control) and the links between nodes reflect which proteins change more similarly in relative abundance across time.

To choose the nodes of the network, we impose two conditions: that the differential expression with respect to the control condition is statistically significant (FDR > 0.05), and that the fold-change in abundance (infected-control ratio) is above a given cut-off, that we chose based on abundance/fold-change plots, Fig. S1A. Whenever these two conditions are met in at least one time point, we select this protein as a node for network construction.

To establish the links between nodes, we need a quantitative measure of ‘similarity’ in abundance changes across time. Since quantified protein levels are always relative abundances (with respect to total counts of ion fragments), usual measures of correlation are biased^73,74^. A better strategy is to interpret co-expression as *proportional* changes across conditions. The idea behind is that if relative abundances of proteins *p_i_* and *p_j_* are proportional across experimental conditions, their absolute abundances must be also in proportion^74^. Proportions are calculated with a log-ratio scaling of protein abundances, and co-expression between every protein pair is quantified by a *proportionality* or *concordance coefficient* ρ_c_ obtained by estimating the log-ratio variances, as^74,75^:

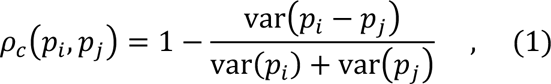

where protein abundances *p_i_, p_j_* have been scaled by a centred log-ratio transformation. This measurement was first proposed as a reproducibility index of independent measurements^76^ and has the same interpretation as a correlation coefficient, ranging between -1 (perfect reciprocity) and +1 (perfect proportionality). To reconstruct the virus-host co-expression network, we consider that two proteins are linked if the absolute value of their concordance coefficient is above a certain cut-off (Table S2). The cut-off value is chosen based on the distribution of concordance coefficients and the number of nodes/links remaining, to avoid either too sparse/disconnected networks or too dense networks (Figs. S1B-D). These values are very similar to the cut-offs defined in other applications of proportionality measures used in the analysis of transcriptomic data^74,75,77^. A recent thorough study on the performance of different association methods to reconstruct cellular networks from scRNA-seq data systematically scored proportionality measures as the best-performing methods^78^.

### 4.3 Network analysis

All network analyses where done in R programming language. We extensively used the *igraph* software package.

#### 4.3.1 Community partition

For community partition and further analyses, we took only the giant component (largest connected component) of the reconstructed network. To identify communities (sets of nodes more densely connected among them than with other sets of nodes) we used the recently developed Leiden algorithm^79^ which is fast and guarantees well-connected communities. Moreover, it provides flexibility in the number of communities defined by changing a resolution parameter γ: higher resolution leads to more communities, while lower resolutions result in fewer communities. To select the best community partition for each virus-host network, we applied the Leiden algorithm to optimize modularity in a range of γ values (0.1 < γ < 2). Simultaneously monitoring the number of different communities and the modularity coefficient, we chose the optimal γ value as the one providing the highest modularity and the most robust community partition (Fig. S2).

#### 4.3.2 Analysis of biological functionality

Functional enrichment or over-representation analysis (ORA) based on Gene Ontology, KEGG and Reactome pathway annotations were conducted for different groups of proteins extracted from the reconstructed co-expression networks (communities, connector nodes and sets of proteins with high both degree and eigenvector centrality). Analyses were undertaken with different functional annotation tools: *gprofiler2*^80^, *clusterProfiler* 4.0^81^ and *GeneCodis4*^82^ with default parameter settings. Statistically significant annotation terms consistently appearing under different pipelines were extracted, and the most relevant annotated proteins manually searched to confirm their biological role. The most significant terms and groups of proteins were represented as dotplots in Fig. 4.

#### 4.3.3. Eigenvector centrality and the theory of interacting networks

The eigenvector centrality can be obtained as the eigenvector **u**_1_ associated with the largest eigenvalue λ_1_ of the adjacency matrix of the network under study^28^. It can be calculated for a single node *i* (as the element *i* of vector **u**_1_), or accumulate it on a set of nodes as the sum of their centralities (normalized such that the total centrality of the whole network is 1). The maximum eigenvalue λ_1_ typically grows with the number of nodes and links of a network, and therefore can be used as a proxy for the strength of a network in a dynamical process where several networks interact. The eigenvector centrality **u**_1_ and the two largest eigenvalues of the system λ_1_ and λ_2_ are directly related to the dynamics as follows: If a process takes place on a network such that **n**(*t*+1) = **M n**(*t*), where **n**(*t*) is the system state at time *t* and **M** is the transition matrix associated with the dynamical system under study, in the limit *t*→∞, **n**(*t*) attains a stationary state independent of the initial condition that coincides with the eigenvector centrality **u**_1_, the maximum eigenvalue λ_1_ yields the growth rate of the population at such stationary state and the time to equilibrium is proportional to ln(λ_1_/λ_2_)^-1^ ^83^.

The dynamical processes that take place on interconnected networks can be analysed as a competition between networks for importance/centrality, measured as the accumulated eigenvector centrality in the nodes of each competitor. An explicit expression can be obtained for the outcome of the competition for centrality between several networks, as well as the time to equilibrium in such competition^25^. A summary of the main results is: (1) the outcome of the competition for centrality between two interconnected networks is only dependent on the largest eigenvalue of the adjacency matrix of each network in the system and on the eigenvector centrality of the connector nodes; (2) increasing the number of interlinks and/or the centrality of the connector nodes of the two competing networks accelerates the dynamical process and increases the final centrality of the weak network, while a low number of interlinks that only connect peripheral nodes will give rise to a very slow process that strongly increases the final centrality of the strong network.

## ACKNOWLEDGMENTS

The authors acknowledge fruitful discussions with I. Pérez-Jover S. Backlund, J. M. Buldú, J. García-Ojalvo and A. Pons. J.A. and R.G. received support from grant No. PID2021-122936NB-I00, J.A. from grant No. MDM-2017-0737 Unidad de Excelencia ’’María de Maeztu’’-Centro de Astrobiología (CSIC-INTA), all of them funded by the Spanish Ministry of Science and Innovation/State Agency of Research MCIN/AEI/10.13039/501100011033 and by “ERDF A way of making Europe”

## AUTHOR CONTRIBUTIONS

J.A. and R.G. conceived and designed the project, contributed to the interpretation of the results and wrote the paper. R.G. performed the numerical analysis of the data.

