## Supporting Information for "Virus-host protein co-expression networks reveal temporal organization and strategies of viral infection"

| | HERPES VIRUSES | | | POX VIRUS |
|---|---|---|---|---|
| | **HSV-1** | **EBV** | **HCMV** | **VACV** |
| **Genome size** | ~152 kbp, > 200 ORFs(*1*) | ~170 kbp, ~100 ORFs(*2*) | ~235 kbp, ~500 ORFs(*3*) | ~190 kbp, >200 ORFs(*4*) |
| **Host cell line** | HaCat (human keratynocite) | P3HR1(Burkitt lymphoma) | HFFF (primary human fetal foreskin fibroblasts) | HFFF (primary human fetal foreskin fibroblasts) |
| **Reference** | Soh et al. 2020(*5*) | Ersing et al. 2017(*6*) | Nightingale et al. 2018(*7*) | Soday et al. 2019(*8*) |
| **# time points** | 7 (0, 2, 4, 6, 9, 12 and 18h pi) | 5(0, 15, 24, 48 and 72h pi) | 8(0, 6, 12, 18, 24, 48, 72 and 96h pi) | 7 (0, 2, 4, 6, 8, 12 and 18h pi) |
| **# viral proteins quantified**[*] | 75 | 64 | 124 | 164 |
| **# host proteins quantified**[*] | 6,875 | 6,791 | 7,635 | 8,271 |
| **# viral proteins differentially expressed (FDR< 0.05)**[#] | 74 | 63 | 119 | 139 |
| **# host proteins differentially expressed (FDR < 0.05)**[#] | 1,261 | 1,209 | 808 | 967 |
| **Immediate early genes/proteins**[&] | $R_{L}2$/ICP0, $R_{S}1$/ICP4, $U_{S}1$/ICP22, $U_{L}54$/ICP27(*9*) | *BZLF1, BRLF1*(*10*) | *UL103, UL104, UL115,UL119*(*11*) | *B11, A48, F11, K1*(*8*) |

**Table S1.** Information on the viruses and data sources re-analyzed in this work. Data were taken from the Supplemental Tables in the provided references. [*]We analyzed only proteins quantified at all time points in biological replicates. [#]Abundance data expressed as pseudo-counts were analyzed for differential expression with respect to the control (non-infected or non-lytic samples) using the *edgeR* pipeline(*12*) and a paired design, controlling for differences between biological replicates. We include proteins differentially expressed with a false detection rate (FDR) < 0.05 at any time point during the course of the infection. [&]Documented immediate early genes of each virus present in the reconstructed co-expression networks, and used for analyses in this work.

| | HERPES VIRUSES | | | POX VIRUS |
|---|---|---|---|---|
| | **HSV-1** | **EBV** | **HCMV** | **VACV** |
| **Absolute fold-change cutoff for number of nodes** | 2 | 2 | 3 | 1.5 |
| **Concordance coefficient cutoff for number of edges** | 0.97 | 0.95 | 0.97 | 0.97 |
| **Number of viral/host nodes giant component** | 71/400 | 59/495 | 76/452 | 134/288 |
| **Number of edges giant component** | 4,430 | 4,840 | 4,521 | 5,323 |
| **Scale-free$** | No | No | No | No |
| **# Communities** | 8 | 6 | 5 | 5 |
| **Modularity coefficient#** | 0.61 | 0.71 | 0.63 | 0.66 |
| **# up/down-regulated nodes** | 74/394 | 278/263 | 144/360 | 137/285 |
| **#positive/negative links** | 4,275/139 | 3,160/1,165 | 3,978/487 | 5,305/18 |

**Table S2.** Parameters and properties of reconstructed virus-host protein co-expression networks. $The scale-free property is assessed by the goodness-of-fit test of the degree distribution to a power law, using a maximum likelihood procedure as described in Clauset et al. (2009)(*13*) and implemented in the *R* package *poweRlaw*(*14*). The modularity coefficient is a measure of the difference between the actual number of edges within communities, and the expected number of edges if they were placed at random, as defined in Clauset et al. (2004)(*15*).

| VACV | # Virus/Host Nodes | Mean deg. | % eigenc. | $\lambda_1$ | $\lambda_2$ | $\lambda_1/\lambda_2$ | # Up/Down nodes | # +/- links |
|---|---|---|---|---|---|---|---|---|
| **Com. 1** | 71/0 | 24.1 | 1.8e-7 | 31.47 | 16.92 | 1.86 | 71/0 | 854/0 |
| **Com. 2** | 62/0 | 30.9 | 2.5e-6 | 35.97 | 14.3 | 2.51 | 62/0 | 959/0 |
| **Com. 3** | 1/13 | 5.6 | 0.03 | 6.74 | 3.21 | 2.10 | 1/13 | 36/3 |
| **Com. 4** | 0/141 | 24.8 | 30.2 | 35.90 | 20.83 | 1.72 | 2/139 | 1,738/12 |
| **Com. 5** | 0/134 | 19.9 | 69.8 | 31.25 | 18.71 | 1.67 | 1/133 | 1,334/2 |
| **Viral net** | 134 | 28.5 | 3.2e-6 | 36.44 | 31.65 | 1.15 | 134/0 | 1,910/0 |
| **Host net** | 288 | 23.7 | 100 | 38.01 | 31.69 | 1.2 | 3/285 | 3,395/14 |
| **EBV** | **# Virus/Host Nodes** | **Mean deg.** | **% eigenc.** | **$\lambda_1$** | **$\lambda_2$** | **$\lambda_1/\lambda_2$** | **# Up/Down nodes** | **# +/- links** |
| **Com. 1** | 59/0 | 30.0 | 94.1 | 35.68 | 11.25 | 3.17 | 59/0 | 885/0 |
| **Com. 2** | 0/13 | 2.5 | 5.9 | 3.16 | 2.37 | 1.33 | 11/2 | 9/7 |
| **Com. 3** | 0/143 | 10.4 | 5.7e-3 | 16.60 | 12.92 | 1.28 | 52/91 | 466/277 |
| **Com. 4** | 0/117 | 14.1 | 0.033 | 25.55 | 12.74 | 2.01 | 71/46 | 416/407 |
| **Com. 5** | 0/107 | 21.6 | 3.3e-3 | 28.92 | 20.51 | 12.36 | 35/72 | 715/438 |
| **Com. 6** | 0/102 | 14.7 | 1.9e-5 | 23.40 | 14.96 | 1.56 | 50/52 | 401/349 |
| **Viral net** | 59 | 30.0 | 94.1 | 35.68 | 11.25 | 3.17 | 59/0 | 885/0 |
| **Host net** | 482 | 15.9 | 5.9 | 31.47 | 25.97 | 1.21 | 219/263 | 2,192/1,633 |
| **HCMV** | **# Virus/Host Nodes** | **Mean deg.** | **% eigenc.** | **$\lambda_1$** | **$\lambda_2$** | **$\lambda_1/\lambda_2$** | **# Up/Down nodes** | **# +/- links** |
| **Com. 1** | 59/52 | 12.7 | 0.3 | 21.24 | 13.57 | 1.57 | 87/24 | 607/98 |
| **Com. 2** | 11/83 | 12.6 | 7.4 | 20.45 | 10.45 | 1.96 | 30/64 | 427/164 |
| **Com. 3** | 2/109 | 12.5 | 4.9 | 19.97 | 13.22 | 1.51 | 22/89 | 591/105 |
| **Com. 4** | 0/73 | 16.4 | 32.8 | 24.23 | 9.82 | 2.47 | 4/69 | 558/42 |
| **Com. 5** | 0/115 | 19.8 | 54.4 | 28.84 | 19.41 | 1.49 | 1/114 | 1,1130/7 |
| **Viral net** | 72 | 8.9 | 0.3 | 16.73 | 8.55 | 1.96 | 72/0 | 319/0 |
| **Host net** | 432 | 17.6 | 99.7 | 32.27 | 27.11 | 1.19 | 72/360 | 3,428/384 |
| **HSV-1** | **# Virus/Host Nodes** | **Mean deg.** | **% eigenc.** | **$\lambda_1$** | **$\lambda_2$** | **$\lambda_1/\lambda_2$** | **# Up/Down nodes** | **# +/- links** |
| **Com. 1** | 68/0 | 11.4 | 0.002 | 16.88 | 12.31 | 1.37 | 68/0 | 389/0 |
| **Com. 2** | 0/25 | 3.4 | 0.003 | 5.06 | 3.31 | 1.53 | 0/25 | 42/0 |
| **Com. 3** | 0/76 | 15.0 | 1.9 | 19.15 | 14.47 | 1.32 | 1/75 | 563/7 |
| **Com. 4** | 0/56 | 16.4 | 11.9 | 21.53 | 9.78 | 2.20 | 0/56 | 458/0 |
| **Com. 5** | 0/62 | 16.8 | 19.8 | 25.06 | 8.04 | 3.11 | 4/58 | 518/3 |
| **Com. 6** | 0/74 | 20.3 | 42.0 | 27.86 | 13.96 | 2.0 | 1/73 | 750/1 |
| **Com. 7** | 0/20 | 5.9 | 0.2 | 7.63 | 4.15 | 1.84 | 0/20 | 59/0 |
| **Com. 8** | 0/87 | 14.1 | 24.2 | 22.45 | 12.48 | 1.80 | 0/87 | 615/0 |
| **Viral net** | 68 | 11.4 | 0.002 | 16.88 | 12.31 | 1.37 | 68/0 | 389/0 |
| **Host net** | 400 | 19.5 | 99.9 | 33.65 | 30.09 | 1.12 | 6/394 | 3,886/11 |

**Table S3. Properties of the different communities and viral/host subnetworks for each virus analyzed**: viral/host nodes per community; mean degree of each community; percentage of eigenvector centrality obtained by community; first and second largest eigenvalues of the adjacency matrix and their ratio; number of upregulated/downregulated nodes per community; number of positive/negative links.

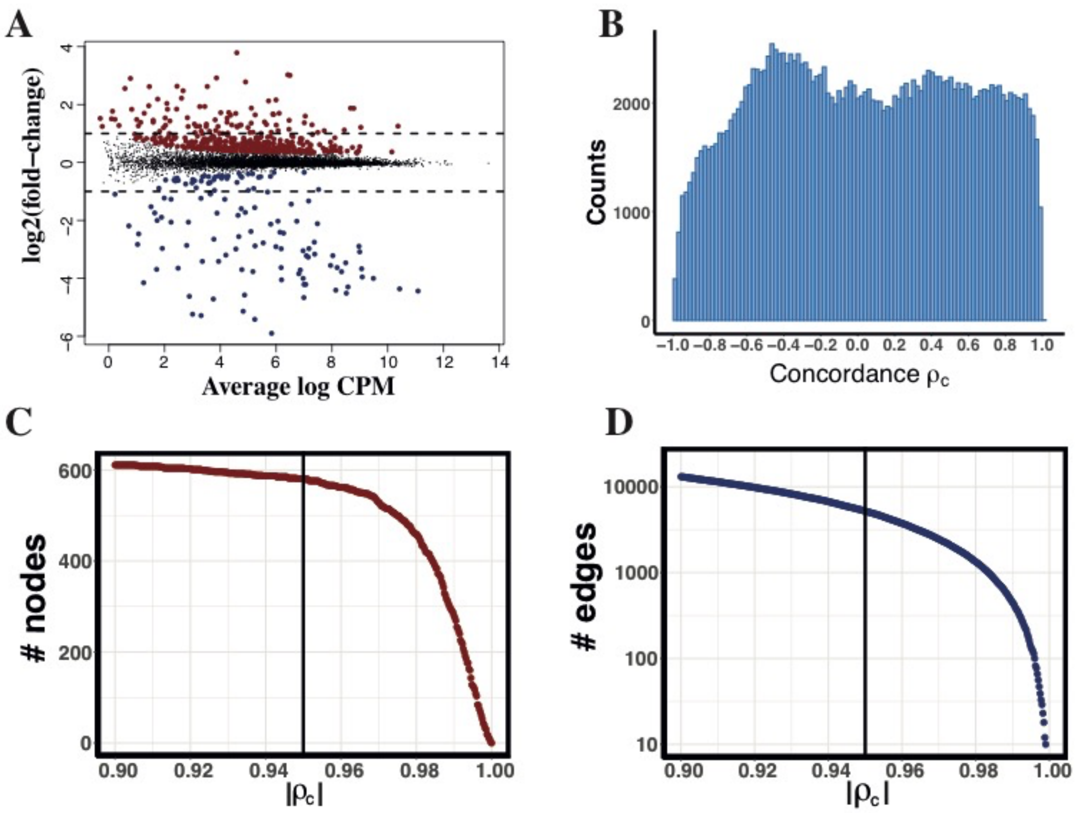


**Figure S1. Choice of cut-off values for network reconstruction.** A. Abundance of each protein (expressed as average over replicates in counts-per-million (CPM)) versus base 2 log of fold-change (relative to uninfected/control samples). Red/blue dots are up/down regulated proteins with FDR < 0.05 (*Methods*). Dashed lines indicate the cut-off value in fold-change to select relevant nodes. B. Distribution of concordance coefficients $\rho_c$ between all selected nodes for the EBV (containing comparable numbers of positive and negative concordances, Table S3). Coefficients are roughly homogeneously distributed between [-0.6,0.9], but frequencies abruptly decrease for $|\rho_c|>0.9$. C. Number of nodes that are left in the network as a function of $|\rho_c|$. D. Number of edges left in the network as a function of $|\rho_c|$. Vertical lines indicate the cut-off value chosen for $|\rho_c|$ giving a good trade-off between number of nodes/edges to avoid too dense or too sparse/disconnected network.

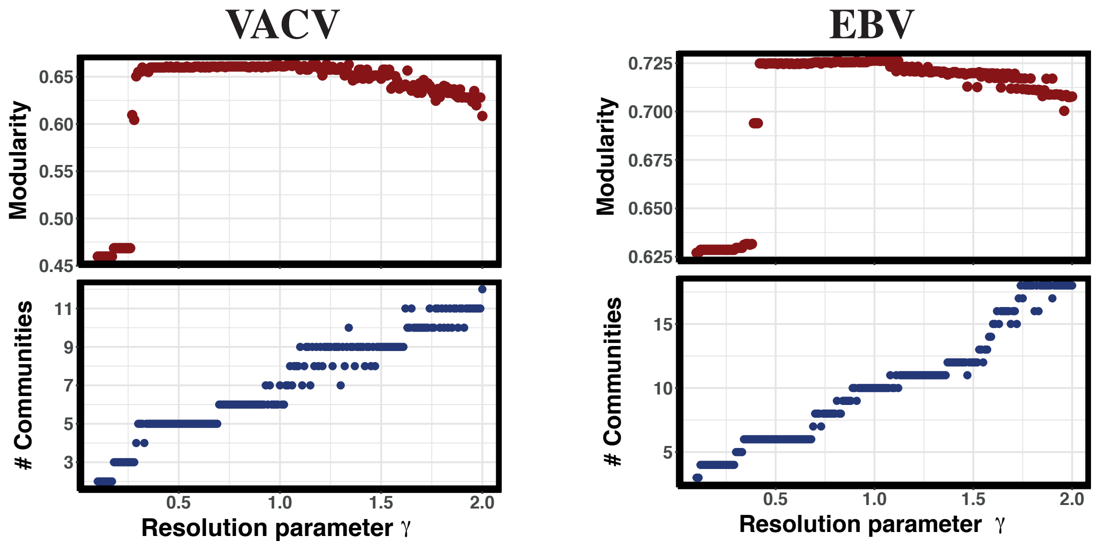


**Figure S2. Selection of resolution parameter for community partition with Leiden algorithm.** Optimized modularity coefficient (top panels) and number of different communities (bottom panels) as a function of the resolution parameter $\gamma$. Taking VACV and EBV virus-host protein co-expression networks as an example, we see that the plateaus in number of communities around $\gamma$=0.5 coincide with a sudden increase in modularity. By inspection of the contingency tables of community memberships of different partitions, we see that plateaus around other $\gamma$ values either result in segregation of few nodes (< 10) from large communities (for instance, the plateau around $\gamma$=0.8 in VACV just adds a new community of 3 nodes split from a large community of 134 nodes) or splits a large community into two, but worsens modularity and/or make partitions less robust (as the plateau around $\gamma$=1.3 in VACV). We thus choose $\gamma$=0.5 as the resolution parameter for these two examples, giving both optimal and more parsimonious and robust partitions.

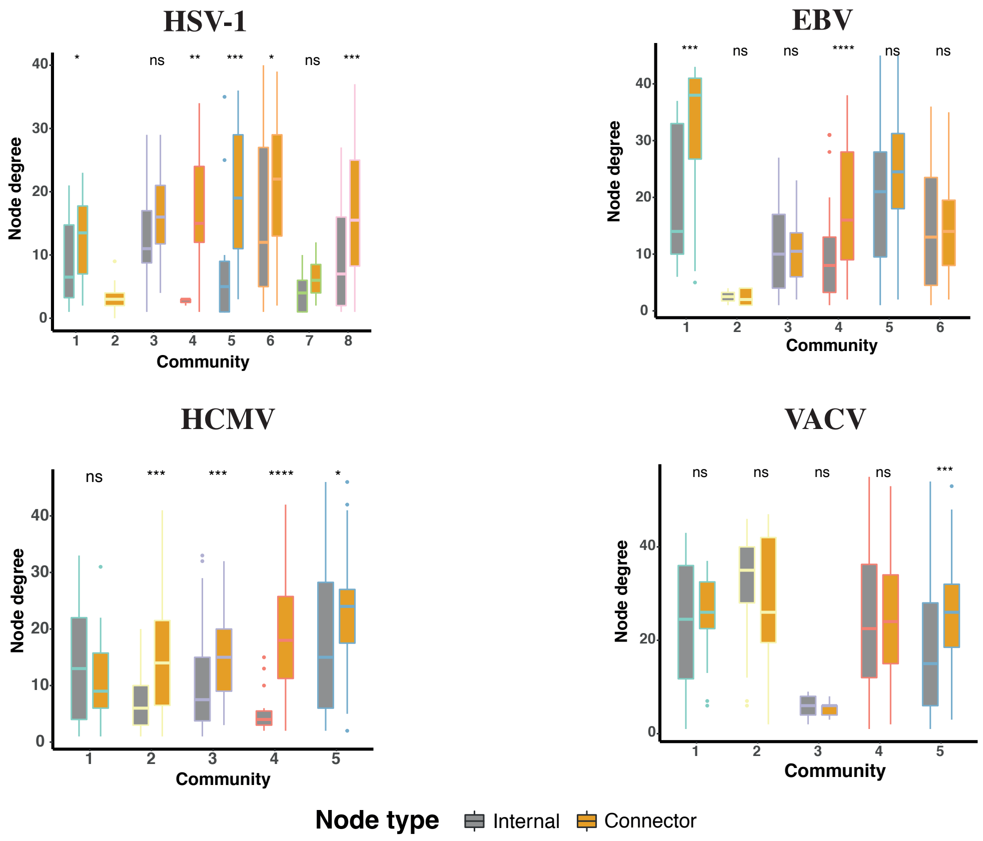


**Figure S3. Degree of internal and connector nodes.**
Distributions of node degrees for each virus type and community, separating internal nodes (nodes connected only to other members of the same community, grey boxes) from connector nodes (nodes connecting to other communities, orange boxes). Significance p-values for difference of the mean between internal and connector nodes are calculated with Wilcoxon test (ns: p>0.05, * : p ≤0.05, ** : p ≤0.01, , *** : p ≤0.001, , **** : p ≤0.0001).

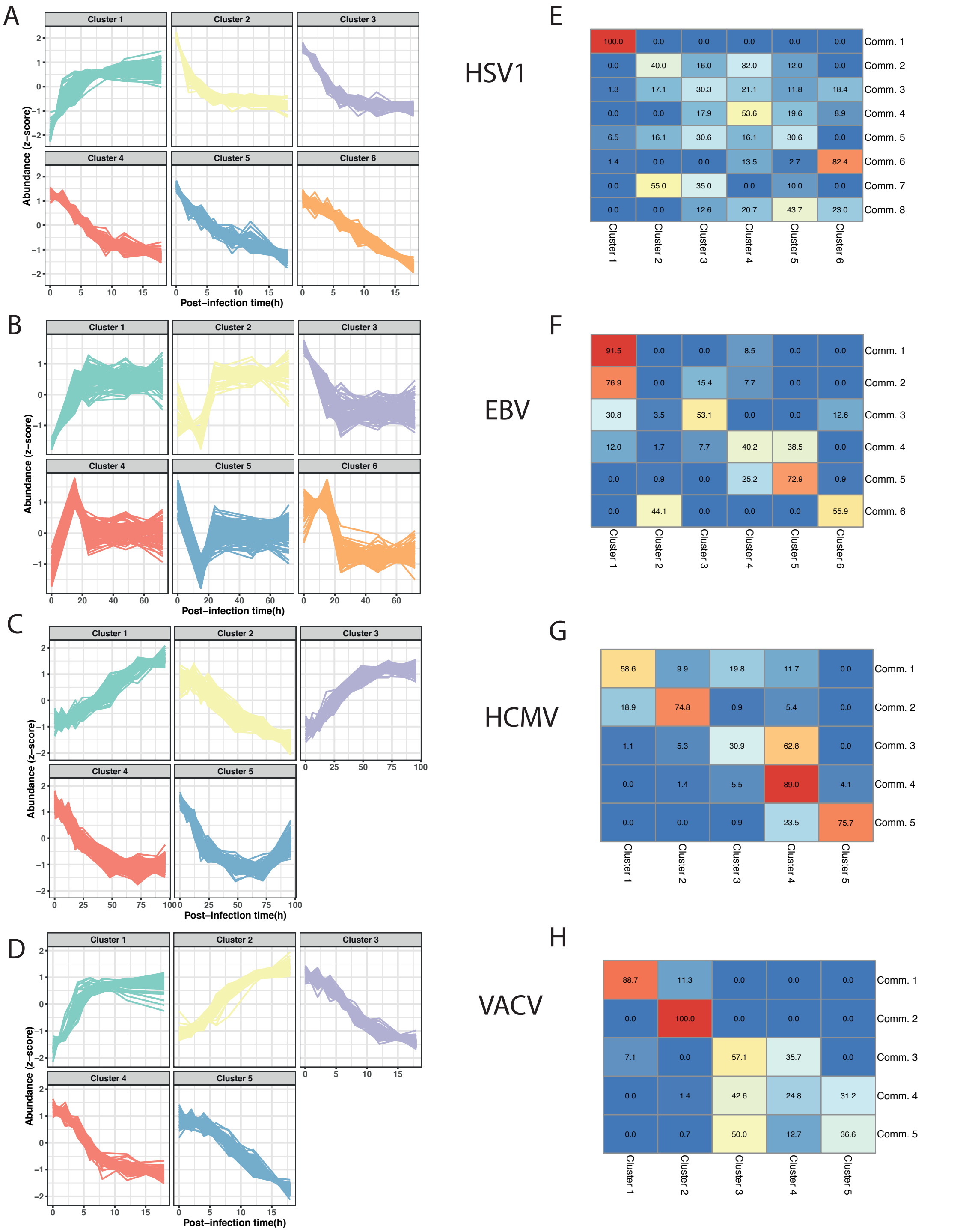


**Figure S4. Clustering of temporal profiles.** Temporal profiles of all network nodes were clustered by similarity using a soft-clustering method employing mixed-effects models and spline fitting(*16*)(‘*TMixClust*’ package in R/Bioconductor). Analysis of silouhette plots(*17*) showed that partitions in 5-6 temporal classes were optimal. A-D: Time series of protein abundances (z-score normalized) grouped by similarity profiles using the clustering method. E-H: Heatmap representation of the contingency table between clustering and community partitions. The numbers represent the percentages of proteins present both in the specified cluster and community.
